# Transcriptional study of the ABC transporter-encoding genes in response to fungicide treatment and during plant infection in the phytopathogenic fungus *Botrytis cinerea*

**DOI:** 10.1101/095869

**Authors:** Elise Loisel, Isabelle R. Goncalves, Nathalie Poussereau, Marie-Claire Grosjean-Cournoyer, François Villalba, Christophe Bruel

## Abstract

The grey mould fungus *Botrytis cinerea* causes worldwide losses of commercially important fruits, vegetables and ornamentals. Various fungicides, with different modes of action, are effective against this pathogen, but isolates with multiple fungicide-resistance phenotypes (Multi Drug Resistance, MDR) have been observed with increasing frequency. In fungi, ATP binding cassette (ABC) transporters participate in drug efflux and we report here on the parallel transcriptional study of the predicted ABC transporter-encoding genes in *B. cinerea.* During plant infection, transcription of all of these genes was activated and over-expression was observed for a few of these genes if the plants were treated with a fungicide prior to infection. In the absence of plant, most of the genes were transcriptionally activated in response to two different fungicides. Both common and specific transcriptional signatures were observed.

## Introduction

*Botrytis cinerea* is the pathogenic fungus responsible for grey mould on grapevines. It is also known for its capacity to infect several other economically important crops, such as tomatoes, raspberries, beans and cucumbers, altogether causing diseases in over 200 plant species [1,2]. Its impact on human activities is, consequently, of acknowledged significance but its direct impact on human health is also gaining recognition. *B. cinerea* could be as important in causing allergies as fungi of the genera *Cladosporium* and *Alternaria* [3,4]. Chemical control has been the method widely used to combat this fungus since the 1950s, and many synthetic fungicides have been intensively used in the field. This, however, has led to the emergence of tolerant strains and *B. cinerea* was one of the first fungus for which resistance was described [5–9].

In microbes that are pathogenic to humans, as well as in human cancer cells, one major tolerance mechanism is multi-drug resistance (MDR), a phenomenon that is usually linked to over-production or increased activity of energy-dependent plasma membrane efflux transporters [10–14]. These transporters belong to two different superfamilies according to the energy source they use to mediate the translocation of molecules. The ATP binding cassette (ABC) transporters hydrolyze ATP and contain a conserved ATP-binding domain [15]. The Major Facilitator Superfamily (MFS) transporters use the proton-motive force to drive transport [16,17]. Members of both families have been shown to exhibit low substrate specificity [18–21].

In fungal plant pathogens, field populations with efflux-based MDR phenotypes have been described essentially in *B. cinerea* [7,22–25] and in a few MDR field strains from *Oculimacula yallundae* [26], *Mycosphaerella graminicola* [27], *Sclerotinia homoeocarpa* [28] and *Zymoseptoria tritici* [29]. In *B. cinerea,* three populations with “MDR phenotypes” have been defined [23]: 1) the MDR1 strains, characterized by a mutation that affects the transcription factor MRR1 and that leads to over-expression of one ABC transporter-encoding gene *(BcAtrB);* 2) the MDR2 strains, characterized by a mutation in the promoter of the *BcMFSM2* gene and its over-expression, and 3) the MDR3 strains, characterized by a genetic recombination between the MDR1 and MDR2 strains that results in an increased expression of *BcAtrB* and *BcMFSM2*. Until now, however, the emergence of these populations has not challenged the current chemical control methods [25,29,30].

Numerous efflux transporters of plant pathogenic fungi are predicted from their genome sequences [31], and their proposed functions are the secretion of virulence factors (secondary metabolites, mycotoxins) and protection against plant defense compounds (phytoalexins), fungicides and molecules produced by various surrounding microorganisms [32–35]. In the genome of *B. cinerea,* Kovalchuk and Driessen have identified 53 putative ABC transporter-encoding genes [31]. The corresponding proteins exhibit different predicted topologies and this has led to their classification into 7 subfamilies (A to G), according to the human genome organization (HUGO)-approved scheme of ABC proteins [36,37].

Four ABC transporters have been characterized in *B. cinerea* (S1 Table). *BcATRA, BcATRD* and *BcATRO* play a role in tolerance to various antifungal molecules, such as cycloheximide, catechol oxopoconazole, fenpiclonil and H_2_O_2_ [38,39]. Their role in the detoxification of natural drugs encountered during plant infection would, however, be limited [37–39]. The fourth, *BcATRB,* plays a role in tolerance to camalexin and resveratrol, two phytoalexins produced, respectively, by *Arabidopsis thaliana* and *Vitis vinifera* [34,40]. Probably as a consequence of this, the Δ*BcATRB* mutant shows a reduced virulence in these two plants. In other plant pathogenic fungi, including, *Fusarium graminearum* [41,42], *Nectria haematococca* [43], *Gibberella pulicaris* [33], *Mycosphaerella graminicola* [44] and *Magnaporthe oryzae* [45–48], ABC transporters also participate in virulence. Orthologues of these transporters are found in *B. cinerea,* but their role in the virulence of this fungus has not been studied (S1 Table).

Multidrug resistance caused by increased ABC transporters activity seems to be regulated at the transcriptional level [23,49,50], but the regulation of these genes is still poorly understood. In addition, some drugs have been shown to affect the transcription of ABC transporter-encoding genes [51–54]. However, in both animal and plant pathogenic fungi where this drug-related regulation has been explored, the studies have been performed in the absence of the fungal host. In this paper we describe the transcriptional behavior of the *B. cinerea* ABC transporter-encoding genes during plant infection, during fungal interaction with fungicide-treated plants, and in response to two different drugs in the absence of plant.

## Materials and Methods

### Fungal strains and growth conditions

The B05.10 strain of *B. cinerea* was grown on malt agar medium (0.5% glucose, 2% malt extract, 0.1% tryptone, casamino-acid and yeast extract, 0.02% RNA, 2% agar) at 21°C under near-UV light. Conidia were collected after 10 days and used to inoculate new plates (5×10^4^/plate), liquid cultures or plants. The other media used for growth were GYPm (1.46% glucose, 0.14% yeast extract and 0.71% peptone) and MSL (0.15% K_2_HPO_4_, 0.2% KH_2_PO_4_, 0.075% (NH_4_)_2_SO_4_, 0.0375% MgSO_4_, 0.375% succinate and 0.15% yeast extract).

### In vitro drug treatment assays

Conidia were collected and calibrated (10^5^/ml) in GYPm or MSL medium. 48-well plates were loaded with 1ml of conidia suspension per well and incubated for 48h, at 21°C, in the dark. Fenhexamid (30μg/ml in DMSO) or Fluopyram (500μg/ml in DMSO) was added to the GYPm and MSL cultures (1μl/well), respectively, while DMSO (1μl/well) was added to the control plates. These concentrations led to a 50% reduction in growth, as measured after 48h by spectrophotometry (OD 620 nm). Mycelia were collected 15, 30 and 60 min after treatment, immediately frozen and lyophilized. Fungicide (technical grade) solutions were freshly prepared from stocks (3g/L) and kept at -20°C. Three biological replicates were generated for both treated and non-treated samples.

### Infection assays

Conidia were collected and calibrated (10^5^/ml) in 0.1% KH_2_PO_4_, 0.2% NH_4_NO_3_, 1.5% gelatin, 5% saccharose. Formulated Fenhexamide (Teldor) and Fluopyram (Luna) were solubilized in water (0.03mg/ml) and sprayed onto cucumber plants (2ml/leaf) 24 hours prior to conidia inoculation; and water was used in the control experiments. Primary leaves from fungicide-treated or non-treated cucumber plants *(Cucumis sativus,* variety “petit vert de Paris”) were inoculated with 20 drops (10 μl) of the conidia suspension. Six primary leaves from treated or non-treated plants were collected after 24 hours (1 dpi), 48 hours (2 dpi), 72 hours (3 dpi) and 144 hours post-inoculation (6 dpi), immediately frozen and lyophilized. All experiments were performed to give three biological replicates.

### RNA extraction and cDNA synthesis

Total RNA was extracted from 4mg of lyophilized ground mycelium or 4mg of lyophilized ground-infected treated or non-treated cucumber leaves using the RNeasy plant midi kit (Qiagen) and according to the manufacturer’s instructions. A DNase treatment (Ambion) was performed to remove traces of genomic DNA. RNA profiles were assessed using the Bioanalyser RNA 6000 Nano kit (Agilent). Double-strand cDNAs were synthesized from 1μg of RNA, using the Thermoscript reverse transcriptase kit (Invitrogen) and according to the manufacturer’s instructions.

### Quantitative PCR and Fluidigm micro-fluidic qPCR

The DNA primers were designed from the cDNA sequence of each gene to be analyzed (S2 Table), using the Primer Express 1.5 software (PE Applied Biosystems), and then synthesized by Eurogentec. qPCR reactions were performed using the Power SYBRH Green PCR Master Mix and an ABI PRISM 7900 HT from Applied Biosystems. Following examination of the primer efficiencies, DNA amplification was carried out as follows: 95°C for 10 min, 95°C for 15 sec and 60°C for 1 min (50 cycles), 95°C for 15 sec, 60°C for 15 sec and 95°C for 15 sec.

Prior to Fluidigm micro-fluidic qPCR analysis, each cDNA sample (1μl) was pre-amplified using a mix of every pair of primers of interest (at a final concentration of 200nM) and the TaqMan PreAmp Master Mix (Invitrogen). Reactions (5μl) were carried out using the recommended program for 14 cycles (10 min at 95°C, 15 sec at 95°C, 4 min at 60°C), diluted 1:5 *(in vitro* assays) or 1:2 *(in planta* assays) and stored at -20°C. The Fluidigm technology reduces qPCR reactions from the usual 10-20 microliter volumes down to 10 nanoliter volumes and permits thousands of DNA amplifications in a single run [55]. Pre-amplified cDNAs and primers were automatically loaded into the chip with the NanoFlexTM 4-IFC controller (Fluidigm). Amplification reactions were carried out in the BioMarkTM real-Time PCR System (Fluidigm) using the following cycling program: 95°C for 10 min, 95°C for 15 sec and 60°C for 1 min (40 cycles). Ct values were extracted from the data using the BioMark Gene Expression Data Analysis software [56].

### Quantification of the transcript levels

A relative quantification of the transcript levels was performed using the comparative Ct method [57]. The PCR signal of the target transcript in the treated sample was compared with that of the control sample at each time point. The relative expression of ABC transporter genes (target genes) to a reference gene was determined using the following formula: relative expression = 2^-∆Ct^ where ∆Ct = (Ct_Target_ – Ct_Reference_). The fold change in expression for a target gene, between a treated sample and a control, was determined using the following formula: fold change = 2^-∆∆Ct^ where ∆∆Ct = (Ct_Target_ – Ct_Reference_)_sample_ - (Ct_Target_ – Ct_Reference_)_control_. The *B. cinerea* actin (accession number BC1G_08198), alpha-tubulin (accession BC1G_02573) and beta-tubulin (accession BC1G_00122) encoding genes were used as references. Three biological replicates were used. The statistical significance of the observed expression differences, between the control and treated conditions, was evaluated with a student t-test on the 2^-∆Ct^ values [57].

## Results

### Expression of the *B. cinerea* ABC transporter-encoding genes during plant infection

After depositing conidia onto cucumber primary leaves, samples were collected prior to primary necrotic lesion appearance (1 day post inoculation (dpi)), after fungal penetration (primary lesions, 2 dpi), during fungal invasion (expanding lesions lesions, 3 dpi) and at the end of the infection cycle when conidiation could be observed (6 dpi) (Fig 1A). Using micro-fluidic dynamic arrays (Fluidigm, [55,57]), expression of the ABC transporter-encoding genes predicted in the genome of *B. cinerea* [31] was studied, in parallel, in a single real-time PCR experiment. Amplification readings could be obtained for 51 ABC-encoding genes out of 53 and primers default likely prevented cDNA amplification for two members of the subfamily C (BC1G_07377 and BC1G_08652). Members of subfamilies E and F encode proteins with no transmembrane domains and which, therefore, cannot be considered as transporters. As a consequence, all the expression data collected for these genes were included in the supplementary materials but are not discussed below.

**Figure 1.**
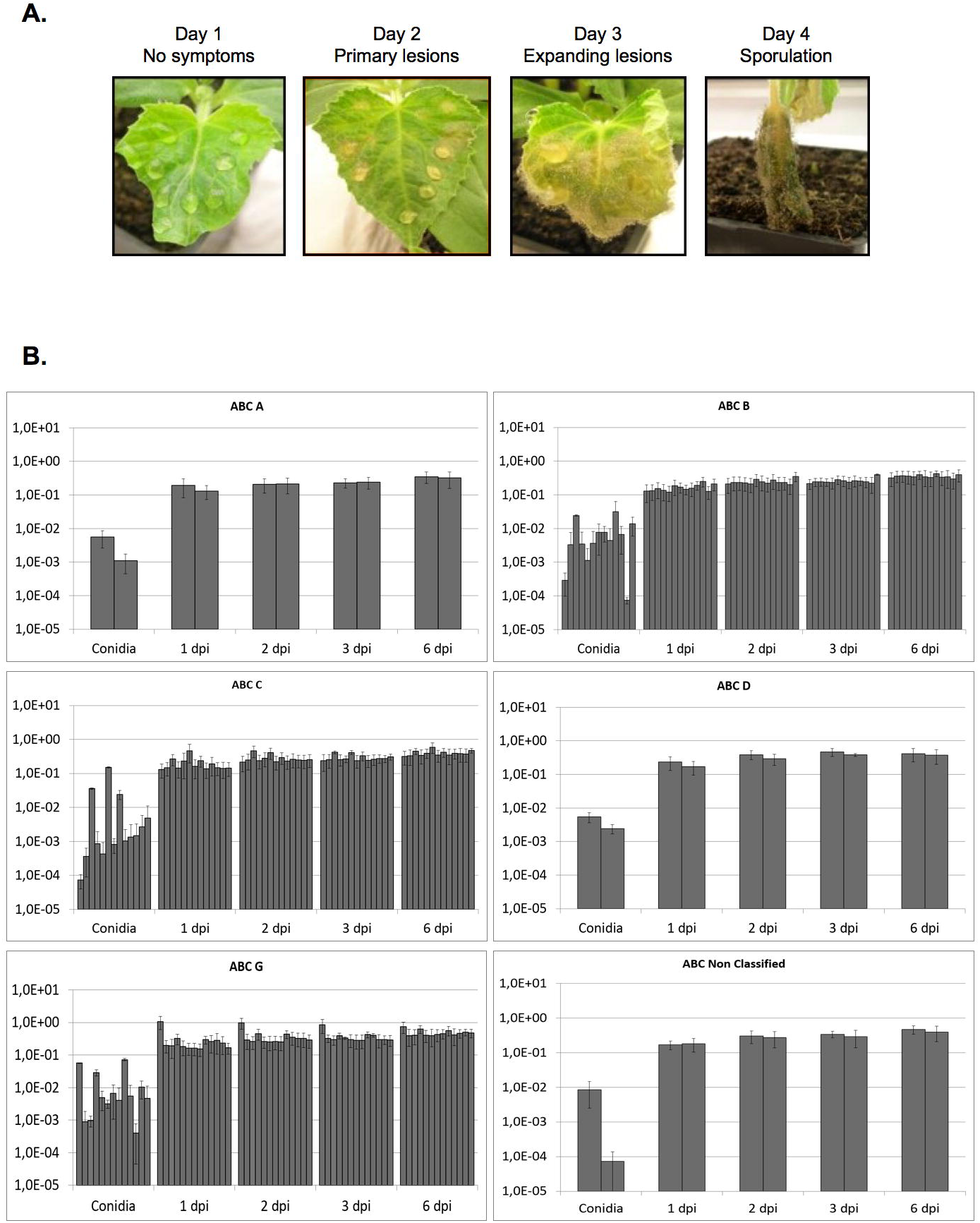
Expression of the *B. cinerea* ABC transporter-encoding genes during cucumber leaf infection. (A) Drops of a calibrated conidia suspension were deposited onto the surface of the leaves. The leaves were harvested at 1, 2, 3 and 6 dpi, immediately frozen in liquid nitrogen and lyophilized prior to RNA extraction. (B) Expression of the ABC transporter-encoding genes inside conidia and the infected leaves (1-6 dpi) were measured by Fluidigm qRT-PCR and relative expressions (2^-∆Ct^) were determined using the comparative Ct method, with the beta-tubulin encoding gene as the reference. Three independent biological replicates were run (standard deviations are shown) and the relative expression are plotted on a logarithmic scale. The subfamilies of ABC-encoding genes are boxed and the expression values are shown for each subfamily member according to its gene number (in increasing order from left to right): **ABC A**; BC1G_11159, 13151. **ABC B**; BC1G_00181, 00244, 02708, 03509, 04774, 05589, 06742, 07740, 10130, 10362, 10568, 10929, 15198. **ABC C**; BC1G_00984, 01001, 01454, 01948, 02001, 06536, 09204, 09469, 09597, 12027, 12625, 14926, 16408. **ABC D**; BC1G_04272, 12469. **ABC G**; BC1G_00425, 01275, 02799, 03332, 04375, 04420, 04465, 05881, 05954, 06867, 08002, 12108, 14379. **ABC NC**; BC1G_03532, 08185. Raw data are available in S3 Table.

In conidia prior to plant contact, as well as on days 1 to 6 of infection, the expression of the reference genes included in the experiment (actin, alpha-tubulin, beta-tubulin) remained unchanged throughout the infection kinetics (data not shown). The beta-tubulin encoding gene was selected to normalize the data because its Ct value was the closest to that of the ABC genes (data not shown). At the conidial stage, variability in gene expression was observed within all ABC subfamilies but this variability was considerably reduced after 1 dpi. In all subfamilies, a significant increase (by a factor of at least 3, and up to more than 2000) in gene expression was observed for all genes between the conidial stage and the infection stages (Fig 1B and S3 Table). Gene expression then either plateaued or increased slightly more (about 2-fold) from 2 dpi to 6 dpi.

We wondered whether the observed expression “jump” between the conidial stage and the first day of infection could be due to the germination process rather than to the beginning of the infection process. To clarify this point, RNA samples that had been used in the transcriptomics analysis of a mycelial plug-driven sunflower cotyledons infection [58] were used to measure the expression of the ABC-encoding genes in a Fluidigm experiment similar to that described above (Fig 2 and S4 Table). Variability in gene expression within the ABC subfamilies was seen in the fungal mycelium used as control in the experiment [58] and considerably decreased after 1 dpi when a general trend of a significant increase in gene expression (by a factor of at least 3, and up to more than 4000) was observed. Gene expression then decreased slightly at 2 dpi, whilst still remaining higher than in the control mycelium. Overall, these results indicate that the transcriptional up-regulation of many ABC-encoding genes at 1 dpi is probably not related to conidia germination, but more likely it is linked to plant infection. These results also show that infection of two different plants, via conidia or mycelium, led to a similar pattern of ABC transporter gene expression.

**Figure 2.**
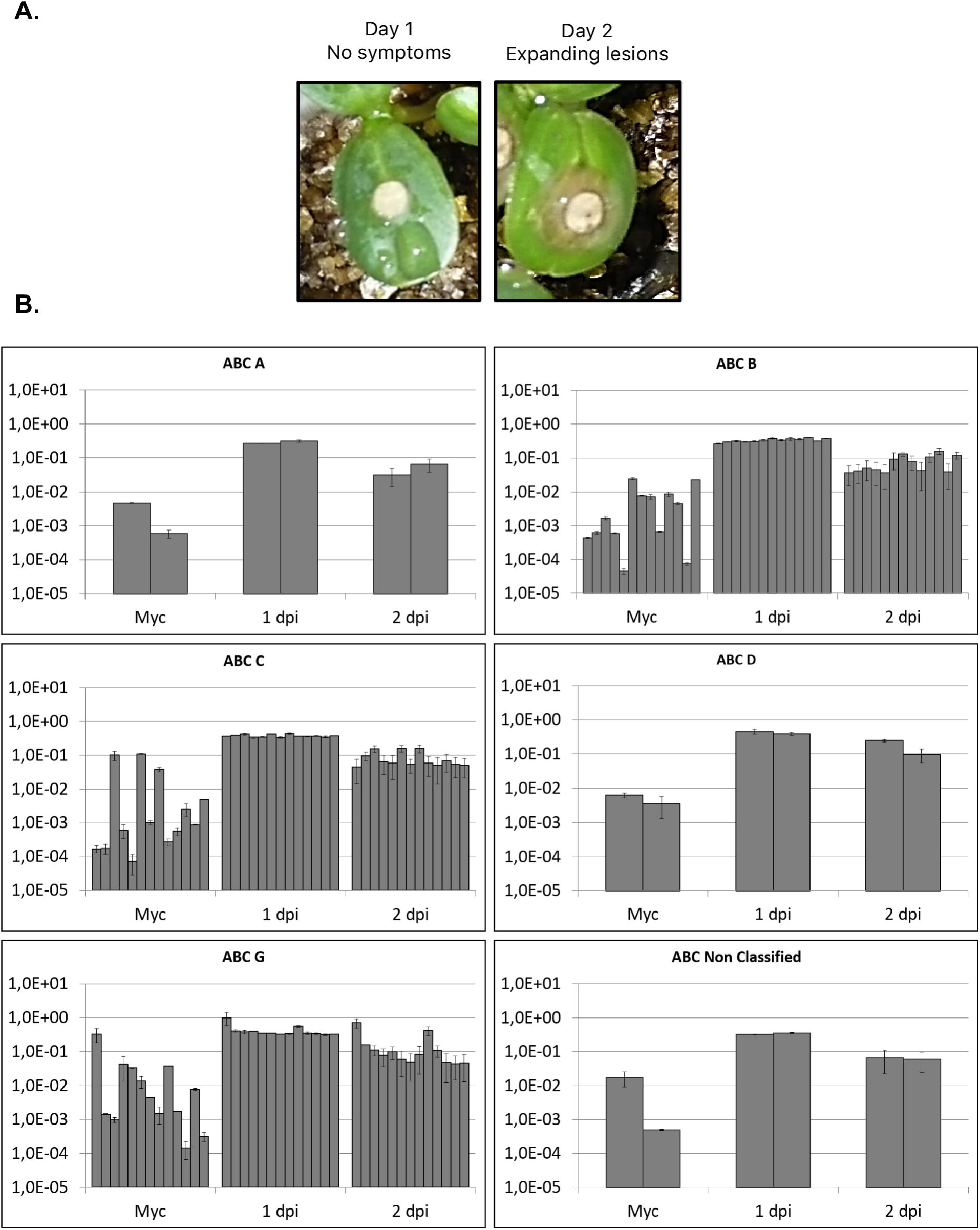
Expression of the *B. cinerea* ABC transporter-encoding genes during sunflower cotyledons infection. (A) A calibrated agar disc from a 48h-old mycelium was deposited onto the surface of sunflower cotyledons. The cotyledons were harvested and immediately frozen in liquid nitrogen at 1 dpi and 2 dpi - (B) RNA extracted from the infected cotyledons and from control fungal mycelium [58] were used to measure the expression of the ABC-encoding genes using Fluidigm qRT-PCR. Relative expressions were determined and plotted as in Fig 1. Raw data are available in S4 Table.

### Expression of the *B. cinerea* ABC transporter-encoding genes during infection of fungicide-treated plants

To study the impact of fungicides on the transcriptional behavior of the ABC-encoding genes during infection, two compounds were selected to treat cucumber leaves prior to fungal inoculation: 1) Fenhexamid, an inhibitor of sterol biosynthesis, used for a long time in the field, and 2) Fluopyram, an inhibitor of Complex II of the mitochondrial respiratory chain [59]. Formulated Fenhexamid and Fluopyram were used to approximate field treatments, and different drug concentrations were tested to establish the conditions of fungal inhibition that would allow reproducibility in the experiments, together with fungal to plant RNA ratios compatible with reliable amplification signals. Concentrations leading to a 25% inhibition of conidia-driven leaf invasion (measured at 3 dpi) were selected for both fungicides. Treated and non-treated cucumber primary leaves were collected at 1, 2, 3 and 6 dpi and RNAs were extracted to run Fluidigm qPCR. In the presence of Fenhexamid, the beta-tubulin-encoding gene showed variations in expression and could not serve as a reference gene. Expression of the actin and alpha-tubulin-encoding genes remained unchanged in the presence of either fungicide, so the actin-encoding gene was used for normalization because it had the closest Ct value to that of our genes of interest (data not shown). Changes in gene expression were observed, at least at one time point, for 22 and 17 ABC-encoding genes when the leaves were treated with Fenhexamid and Fluopyram, respectively, and 16 of these genes were activated in response to both chemicals (Fig 3 and S5 Table). In response to Fenhexamid, the expression of most genes increased at both 1 dpi (21 genes out of 22; mean fold change = 5.6) and 6 dpi (12/22; mean fold change = 3.1). In response to Fluopyram, the expression of most genes (13 genes out of 17) increased from 1 dpi to 3 dpi (with a peak in expression at 3 dpi; mean fold change = 5.1).

**Figure 3.**
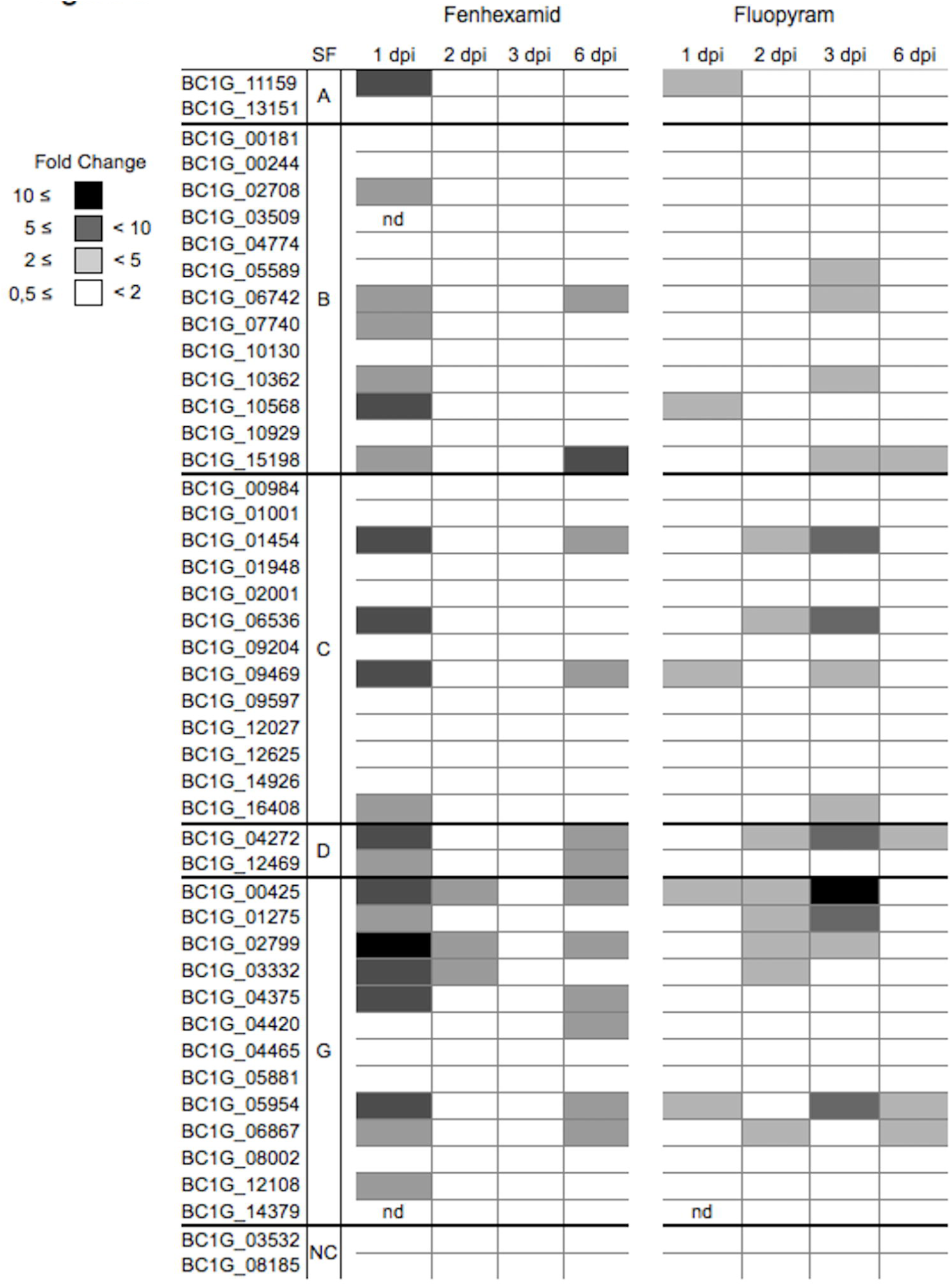
Expression variations of the *B. cinerea* ABC transporter-encoding genes in response to fungicide treatments during cucumber leaves infection. Non-treated and Fenhexamid or Fluopyram-treated primary cucumber leaves (treatment 24h before infection and concentrations set to reduce leaf invasion by 25% at 3 dpi) were infected using conidia and then collected, over six days, to extract RNA. Gene expressions were measured using Fluidigm qRT-PCR with the actin encoding gene as the reference (see S5 Table). Fold changes were determined between the treated and the untreated sample collected at the same time point. Three independent biological replicates were run. The ABC transporter-encoding gene subfamilies (SF) are indicated by their one letter code or NC (Non-classified). Expression fold change values are represented with a grey scale defined on the figure. “nd” stands for no amplification read for the corresponding gene in this experiment.

### Expression of the *B. cinerea* ABC transporter-encoding genes in response to fungicides in the absence of plant

A fast transcriptional response (30 min) of ABC-encoding genes to fungicides has been observed [25,40,60], but this could not be measured in the experiments presented above due to the difficulty of collecting sufficient amounts of fungal material (and, therefore, extracted RNA) after very short periods post inoculation of the plants. Consequently, some additional experiments were set up in the absence of plant to analyze the behavior of the ABC-encoding genes shortly after the exposure of *B. cinerea* to Fenhexamid and Fluopyram. Drug concentrations that reduce fungal growth by 50% were selected to match the conditions used in other similar studies [39,60].

Young mycelium, grown 48h *in vitro* in the absence of any drugs, was exposed to Fenhexamid for 15, 30 or 60 minutes and total RNA was extracted from the mycelia collected at each time point to run Fluidigm qPCR. Thirty two genes out of 45 showed higher expression in the treated samples than in the control untreated samples, at least at one time point (Fig 4 and S6 Table). Nine genes were over-expressed by more than 10-fold in response to the fungicide, 7 showed a 5 to 10-fold over-expression and 16 showed a 2 to 5-fold over-expression. The remaining 13 genes were not differentially expressed. The vast majority of members of the ABC-B (92%; 12/13) and ABC-C (77%; 10/13) subfamilies showed significantly elevated transcript levels, as did the two ABC-D genes. In all the other ABC subfamilies, one out of two genes was differentially expressed; 1/2 ABC-A, 6/13 ABC-G and 1/2 NC. These results indicate that *B. cinerea* responds quickly to Fenhexamid by the transcriptional modulation of a large number of ABC transporter-encoding genes.

**Figure 4.**
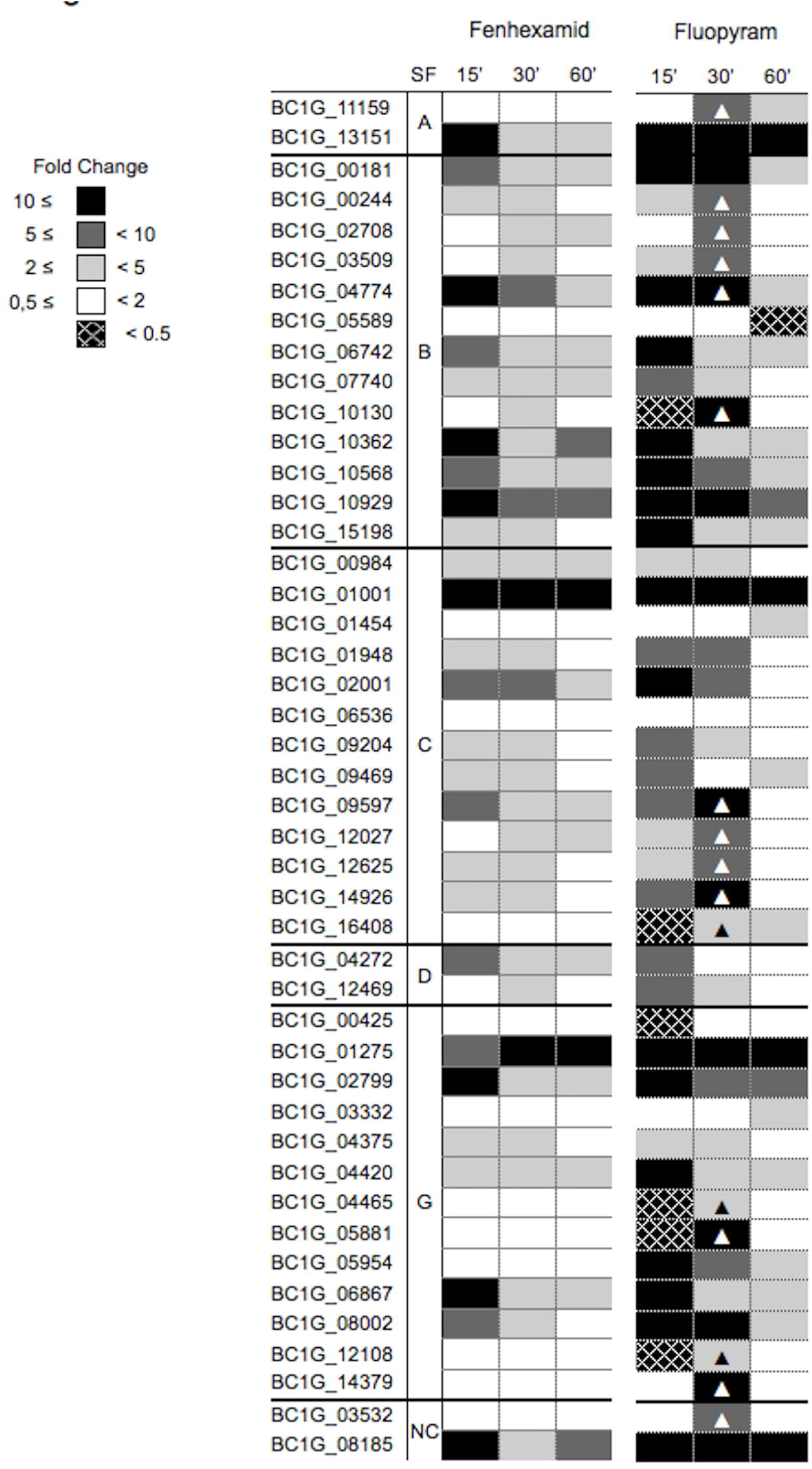
Expression variations of the *B. cinerea* ABC transporter-encoding genes in response to fungicide treatments in the absence of plant. Fungal mycelium grown 48h in the absence of plant and in the absence of fungicide was then exposed to Fenhexamid or Fluopyram (concentrations set to reduce growth by 50%) for 15, 30 and 60 min. Expression fold change values were obtained as in Fig 3. Triangles highlight the genes whose expression was maximum at 30 min post-treatment. Raw data are available in S6 Table.

Under the same experimental conditions described above for Fenhexamid, we observed that Fluopyram also caused wide transcriptional modulation of the ABC transporter-encoding genes (Fig 4 and S6 Table). In comparison with the Fenhexamid experiment, the transcriptional response was even more pronounced since 44 genes out of 45 (98%) were differentially expressed. Moreover, 22 genes were up-regulated by more than 10-fold, compared with only 9 genes in response to Fenhexamid, and 13 genes were up-regulated by between 5 and 10-fold, again compared with only 7 genes in response to Fenhexamid. All of the genes (27) up-regulated 15 minutes after Fenhexamid exposure were also up-regulated 15 minutes after Fluopyram exposure, and a strong positive correlation (Spearman’s rank analysis; rho=0.88; p-value=2.2×10^-16^) could be observed in the expression data collected after these treatments (S1 Fig). This result shows that two different fungicides trigger a similarly fast and wide transcriptional response among the ABC-encoding genes in *B. cinerea.*

At last, sixteen genes showed a maximum up-regulation at 30 minutes after Fluopyram exposure (fold change at 60 minutes decreased) (Fig 4). This peak of up-regulation in response to Fluopyram suggested a specific transcriptional response for this group of genes.

## Discussion

The expression of the putative ABC transporter-encoding genes in the phytopathogenic fungus *B. cinerea* was investigated throughout the course of plant infection, during the infection of fungicide-treated plants and in response to fungicide exposure *in vitro.* Two different fungicides were tested - one inhibiting sterol biosynthesis (Fenhexamid) and one inhibiting the respiratory chain Complex II (Fluopyram). Using high throughput Fluidigm technology, the parallel expression of 45 genes, classified in subfamilies A, B, C, D, G, and “non-classified”, could be measured.

### ABC transporter transcriptional response during plant infection

Under our experimental conditions, the expression of all the ABC transporter-encoding genes showed an “expression jump” on the first day of infection when cucumber leaves were infected using conidia or when sunflower cotyledons were infected using mycelium plugs. This may indicate that efflux plays a role at an early stage of pathogenesis, before symptoms are visible. Transcriptional activation of the ABC-encoding genes could be related to detoxification and/or the secretion of virulence factors [44,61]. Indeed, toxic compounds, such as phytoanticipins and phytoalexins, are released by plants following an attack by a pathogen [62] and some of them are known to induce the expression of transporter-encoding genes in *B. cinerea* [34,39,40], *N. haematococca* [43] and *M. oryzae* [47]. It is also acknowledged that fungal ABC transporters can protect the fungal cell against the intracellular toxic accumulation of biomolecules that they have produced [46] and that they can secrete secondary metabolites [63]. In *B. cinerea,* almost 90% of the genes involved in the biosynthesis of secondary metabolites belong to probable co-transcribed DNA clusters, and 40% of the latter contain one ABC or MFS transporter-encoding gene [58]. The increased expression of ABC-encoding genes during plant infection could, therefore, be correlated with the secretion of such metabolites. Finally, a few ABC transporters have been shown to play a role in virulence (S1 Table) and all the corresponding genes in *B. cinerea* are induced during infection of cucumber and sunflower leaves. The expression level of all genes remained high over a 6-day infection period on cucumber leaves and this may indicate that efflux plays a long-term role during pathogenesis. In the rice pathogen *M. oryzae,* over-expression of all ABC-C encoding genes has been recorded during the early and late phases of plant infection [52], but in other studies on *B. cinerea,* this was not observed, either for *BcatrA,* whose activation was reported only during the first day of bean leaf infection [60] or for *BcatrO,* whose transcript accumulation peaked 6h after fungal inoculation [39].

### ABC transporter transcriptional response to fungicides in planta

When considering crop protection, information related to the transcriptional response of ABC-encoding genes to fungicides in the context of an infection seems to be of interest. A correlation between an increased expression of a few ABC-encoding genes (*atrA, atrB, atrK* and *BMR3*) and MDR phenotypes has been found in *B. cinerea* field strains [7,23,64] and over-expression of ABC-encoding genes could be expected in response to fungal contact with treated plants. Here we show that several ABC-encoding genes were induced in *B. cinerea* in response to Fenhexamid or Fluopyram *in planta.* However, a small difference in transcript levels was observed when compared to the level of gene induction that was recorded during infection of non-treated plants. This could relate to the low dosage of fungicides applied onto the leaves or to a near-maximum level of expression achieved onto non-treated leaves. We attempted to clarify this issue by using higher doses of fungicides that inhibited 50% and 75% of conidia-driven leaf invasion instead of 25% (measured at 3 dpi), but the experiments were not reproducible under these conditions.

Most genes (75%, 15/20) whose expression was induced in response to Fenhexamid *in planta* were also induced in response to Fluopyram. The expression profiles were, however, different over the course of infection. In Fenhexamid-treated leaves, transcriptional induction occurred during the asymptomatic stage (1 dpi) and during the conidiation stage (6 dpi) while it peaked during the colonization stage (2 dpi and 3 dpi) in Fluopyram-treated leaves. These profiles might relate to changes in drug concentration at the cellular level, to the relative importance of the targeted fungal processes during infection, or to changes in the physiological state of the fungus, as previously suggested by Judelson and Senthil [51] in *P. infestans.* Lastly, we note that the well-studied *BcatrB* gene responded only to the presence of Fenhexamid and this supplements what is currently known about this gene and demonstrates that its expression is not induced in response to all fungicides.

### ABC transporter transcriptional response to fungicides in the absence of plant

In the absence of plant, the expression of 69% (31/45) of the ABC transporter-encoding genes was activated 15 minutes after exposure to either Fenhexamid or Fluopyram. A similar rapid response has already been reported in *B. cinerea* for the *BcatrA* gene (BC1G_03332), whose activation started 10 minutes after fungal exposure to cycloheximide and reached a maximum (fold change of about ten) 50 minutes later [60]. A rapid response was also reported for the atrB gene after treatment of *A. nidulans* with the azole fungicide fenarimol [25]. Our results therefore support the observed capacity of fungi to rapidly detect and react to xenobiotics, or to the cellular disturbance that they induce.

Twenty seven genes were up-regulated both 15 minutes after exposure to Fenhexamid and 15 minutes after exposure to Fluopyram. Moreover a positive correlation was detected between the transcriptional responses induced by the two fungicides. This result suggests that a large number of the same ABC transporters are produced by *B. cinerea* to adapt to the presence of different toxic molecules. A general stress response to growth inhibition (50% in this study) or a more specific response to fungicides could account for the result, but in both cases this reveals a fast "broad" response to poisoning that might represent a first line of defense against multiple xenobiotics. The production of many transporters, each of them able to efflux more than one chemical, could efficiently and rapidly protect the cell from a large number of molecules. In accordance with this hypothesis, Kim *et al.* have shown that 3 different fungicides induce the transcription of 9 to 11 ABC-C members in the rice pathogen *M. oryzae* [52]. Other studies have also shown that a substrate to one ABC transporter can lead to both the transcriptional activation of the corresponding gene and to that of one or more other ABC transporter-encoding genes belonging to different subfamilies [34,48,51,65]. Finally, the possible activation of multiple yeast ABC transporter genes (of different subfamilies) by one transcription factor has been proposed (PDR network [49,66,67]). In *B. cinerea,* the transcription factor MRR1 (Multidrug Resistance Regulator 1 [23]) could play a role in the "broad" response described above since the group of 27 genes concerned (up-regulated after exposure to Fenhexamid and Fluopyram) contains its known target *BcatrB.* On the other hand, no clear homologue of the yeast PDR regulators has been identified in *B. cinerea* and the yeast nuclear receptor-like induction mechanism of ABC-encoding genes [68–72] is unlikely to be operational in *B. cinerea* and to play a role in the “broad” response.

The expression of 9 genes was activated after exposure to Fluopyram and not after exposure to Fenexhamid. Furthermore, the expression of 16 genes showed a maximum expression 30 minutes after exposure to Fluopyram and not after exposure to Fenhexamid. Lastly, iprodione and resveratrol induced the expression of *BMR1* (BC1G_00425) in *B. cinerea* [73] while no change was recorded in the expression of this gene in this study. Overall these results indicate that *B. cinerea* is able to react differently to different drugs, in addition to the “broad” response described above.

In conclusion, our study revealed a transcriptional up-regulation of a large set of ABC transporter-encoding genes during the interaction of *B. cinerea* with a plant. Following fungicide-plant treatment, additional but moderate up-regulation was recorded for multiple genes and two different expression profiles were observed over the course of the infection of plants treated with two different drugs. Lastly, parallel *in vitro* experiments revealed a possible fast and “broad” fungal response to xenobiotics. Molecular understanding of this response and testing its existence in other fungal pathogens could be of interest since it might impact fungicide treatments.

## Acknowledgments

We are grateful to A. Rolland, T. Van Der Merlen and D. Paelink from Bayer-SAS for their technical assistance in the Fluidigm experiment. We thank G. Mey for the initiation of the project, C. Rascle for technical support and M. Choquer for discussions. This work was supported by the CNRS, the University Claude Bernard Lyon 1 and Bayer SAS.

## Supporting Information

**S1 Fig. Correlation between expression fold changes.** Expression fold changes observed in the absence of plant between untreated samples and samples exposed to either Fenhexamid or Fluopyram during 15 min are correlated (S6 Table).

**S1 Table. ABC transporters genes IDs correspondence in *B. cinerea.***

**S2 Table. Primers used in the study**.

**S3 Table. ABC-transporter gene expression during cucumber leaf infection**.

**S4 Table. ABC-transporter gene expression during sunflower leaf infection**.

**S5 Table. ABC-transporter gene expression in response to fungicide treatments during cucumber leaf infection**.

**S6 Table. ABC-transporter gene expression in response to Fenhexamid treatments in the absence of plant**.

**S1 File. Upstream sequences, in fasta format, of ABC-transporter genes showing a maximum expression at 30 minutes after Fluopyram exposure**.

